# Endogenous gene tagging with FnCas9 to track and sort neural lineages from 3D cortical organoids

**DOI:** 10.1101/2025.01.09.631995

**Authors:** Manoj Kumar, Berta Terre Torras, François Guillemot, Debojyoti Chakraborty

## Abstract

Neural lineage tracing, or molecular dissection of lineage-specific brain cell types, is used in many labs to learn how neurons grow and mature. However, these studies depend on the growth and characterization of pure cultures, which takes a long time because of biochemical or fluorescence-based isolation through cell surface markers that overlap. These lineage-specific cells, however, can be efficiently sorted using endogenously expressed, fluorescently labeled marker genes. The labeled cell lines can be used not only to differentiate and purify different types of neurons but also to study the long-term development of neural lineages in two- and three-dimensional development models. In this study, we used an orthogonal Cas protein to generate human embryonic stem cell (hESC) lines with genetically labeled fluorescent barcodes for discrete neural lineages. We use these lines to successfully demonstrate spatial and temporal tracing of DCX-positive neuroblasts and immature neuronal cells within 2D neural cultures and 3D cortical organoids derived from human embryonic stem cells. This allowed the purification of endogenously tagged live neural cells from heterogeneous cortical organoids across multiple stages of development.

## Introduction

In an effort to comprehend brain functions, approximately 25 cell classes—16 neuronal and 9 non-neuronal—have been identified in the human brain to date (1–3). However, by examining gene expression in single brain cells, researchers have discovered potentially significant cell types in about 100 clusters (4–13). The complexity of the human brain is due to the many different types of neuronal cells that grow from the neural stem cells very early on during development and the spatial and temporal expression of genes that regulate their growth and proliferation (14–16). To date, many studies have been performed to study these neural subtypes in depth for their roles and to understand their complex interconnections (3, 17–22). In the last decade, advances in cell biology have made it possible to grow or obtain these neural lineages in the lab by differentiating embryonic stem cells or neural stem cells in both 2D and 3D cultures (23–34). However, while studying them as a single population, it is hard to dissect the different cell lineages because of their origin from single neuroepithelial cells and the abundance of identical surface markers in several cases (35–39). Thus, the purification of lineage-specific cells based on traditional biochemical or fluorescence-based surface markers is challenging. Furthermore, other reported neural lineage tracing methods are not designed for lineage-specific, in-depth molecular analysis of pathways that regulate neuronal development, and methods for tracing the lineage of a cell based on scRNA-seq often do not include spatial information, rendering reconstruction of temporal expression data challenging (40–42).

As an alternative, the expression of endogenous lineage-specific marker genes has the ability to identify and isolate such cells efficiently by linking expression with cell type and time in development. For example, isolating intermediate progenitor cells would require endogenous tagging of TBR2 (T-box brain protein 2, also known as Eomesodermin) (43). Similarly, neuroblasts, or immature neuronal cells, can be isolated using DCX (doublecortin) expression (44). Therefore, endogenous marker genes tagged with fluorescent reporters can be used to efficiently isolate neural subtypes with greater purity (42, 45, 46). To meet the need for marker-specific neural lineage tracing *in vivo*, we propose CRISPR/Cas9-mediated endogenous marker-specific gene-tagging in 3D organoid models for marker-specific lineage tracing while preserving cell identity and spatial existence. Although such approaches have been explored in the past in the context of embryonic stem cells and certain differentiation markers in pure cultures, their adaptability in 3D or organotypic cultures has been relatively unexplored.

CRISPR/Cas9’s ability to make double-stranded DNA breaks (DSBs) at the desired target locus and trigger HDR (homology-directed repair)-mediated DNA knock-in has been used in a number of ways, from fixing mutations in DNA to adding reporter systems (47–53). With the help of CRISPR/Cas9, it is possible to insert a gene that codes for fluorescent protein at a genomic locus *in vivo* (52–55). This can be used for endogenous gene tagging, real-time protein dynamic assays, and other ways to track proteins (52–57). However, when such large HDR-mediated DNA knock-ins are performed, the efficiency of the knock-in is inversely related to the length of the DNA template (58). As highlighted in several of our previous reports, FnCas9, a Cas9 system isolated from the bacterium *Francisella novicida*, can introduce such single-large donor DNA templates much more efficiently as compared to the conical SpCas9 (59, 60).

With the help of FnCas9 and some experimental optimization, we now show that in hESCs, endogenously tagged marker genes can be efficiently used for spatio-temporal expression of lineage markers, leading to the successful dissection of cell types. As a proof of concept, we showed that when these hESCs were differentiated into 2D cortical cultures, the stage-specific expression of DCX-tdTomato could be seen. This confirmed the development of a 2D neural differentiation model for endogenous marker-specific neural lineage tracing. We could also validate these endogenously tagged *DCX*-tdTomato hESCs by differentiating them into human cortical organoids (hCOs) and showing the co-existence of DCX as well as the tdTomato fluorophore, validating the proper expression of the genetically encoded reporter. Further, due to the biological activity of tdTomato fluorophore, we were finally able to show that DCX-positive neuronal cells could be easily traced and extracted from 3D cortical organoids by fluorescence-assisted cell sorting (FACS) in a mixed cell population. Collectively, we show an end-to-end optimization of HDR-mediated single-large knock-in in human embryonic stem cells (hESCs) to endogenously label DCX-tdTomato+ neuroblast cells in developmentally related 2D and 3D models for neural lineage tracing.

## Results

### FnCas9-based HDR-mediated knock-in of a single-large DNA template

The HDR-mediated DNA template knock-in depends on a number of factors, including but not limited to the site of integration, nature of the donor construct, length of the homology arms, etc. (61–63). In the past, several studies have demonstrated successful Cas9-based HDR-mediated integration of large DNA for downstream applications. However such integrations are hard to do in different types of cells, and HDR-mediated knock-ins of large donor DNA templates tend to have low integration rates (61). In our earlier studies, we have demonstrated that the orthogonal Cas9 from *Francisella novicida* (FnCas9) outperforms SpCas9 for HDR-mediated knock-in of donor constructs (59, 60).

We tested the ability of FnCas9 to perform a similar knock-in in h9 hESCs. and used PCR genotyping to see if the reporter construct was successfully integrated (Figure 1). We designed a single donor DNA template with a tdTomato gene in the frame and a P2A (self-cleaving peptide) sequence in the left homology arm. P2A was introduced to stop the formation of a recombinant DCX protein that can cause functional interference with the endogenous protein while fluorescently labeling cells that express DCX. To simplify the process of selecting positive knock-in events without the need for colony picking or cell sorting, we also included an antibiotic resistance gene with an independent pGK promoter flanked by LoxP sites in the construct. Once successful integrants are found, LoxP sites would allow the antibiotic gene to be cut out when treated with Cre, and expanded sites would get rid of it when the positive knock-in cells were chosen over the right antibiotic (54, 65). The donor DNA construct also contains 300 bp of homology arms on each side of the left and right homology arms. These arms have the carrying sequences from the endogenous DCX C-terminus that the designed gRNA is meant to target.

**Figure 1:**
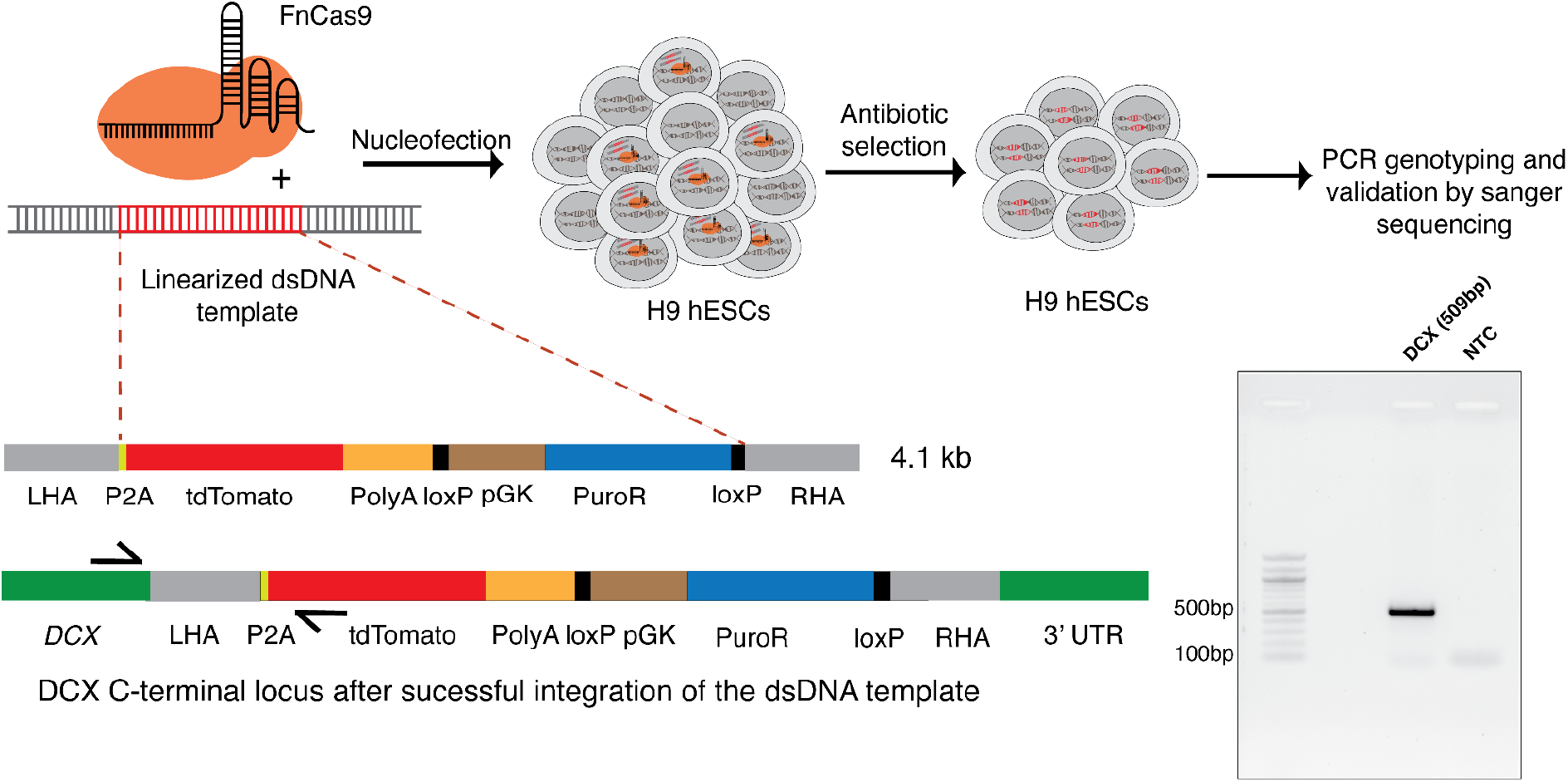
FnCas9-based HDR-mediated knock-in of a single large DNA template; schematic of the process that was optimized for FnCas9-based HDR-mediated endogenous tagging of the selected marker gene, DCX. The linearized donor DNA template used for knock-in is shown with various components, along with left and right homology arms complementary to the C-terminus end of the *DCX*. The agarose gel shows PCR genotyping of the antibiotic-selected cells with the successful integration of the DNA donor template. (NTC, no DNA template control)

We were able to successfully isolate stem cell colonies where the DCX-tdTomato donor DNA template is integrated into the genomic DNA of the selected h9 hESC population, as confirmed by PCR-based genotyping. (Figure 1). This was further validated by performing Sanger sequencing and comparing the expected DNA sequence to the results obtained (Supp. Figure 1a). To use this strategy on other similar loci, we added an endogenous tag to the C-terminus of the intermediate progenitor cell marker TBR2 and used PCR genotyping (Supp. Figure 1b) to confirm the integration. As iPSCs do not show expression of either DCX or TBR2, we next proceeded to validate the functional expression of the reporter fluorophores in the expected neural cell types by differentiating the iPSCs into 2D neural cultures or 3D cortical organoids.

### 2D cortical differentiation of FnCas9 gene-tagged h9 hESCs to track neural lineages with specific endogenous markers

Due to the pluripotent nature of embryonic stem cells, genetically manipulating them allows the dissection of pathways that appear further downstream in developmental stages. Human embryonic stem cells (hESCs) have been successfully differentiated towards cortical neurons with the help of a previously optimized cocktail of a few SMAD inhibitors (66–68). Using these published protocols to direct hESCs to become cortical neurons, we adapted the protocol depicted in Figure 2a as well as the methods (69). We followed a similar approach with slight modifications (the details are in the methods section). After treating DCX-tdTomato hESCs with dual SMAD inhibitors like dorsomorphin and SB-431542, these cells were pushed towards the initiation of neural differentiation and kept in cultures to look for fluorescence signals. Around day 23 of the 2D neural differentiation protocol, the cells appear to be forming rosette-like structures, with tdTomato expression becoming visible (Figure 2b). Following further maintenance in culture, these cells exhibited the morphology of early neuroblasts through the presence of neurites and a strong expression of endogenously tagged DCX-tdTomato, making them appear bright red (Figure 2b). Keeping the cells in culture until day 53 allowed the neural processes to reach maturity, becoming longer and showing prominent branching morphology. The continuous expression of tdTomato suggested that the cells were neuronal precursors or immature neurons. Importantly, DCX-tdTomato expression throughout the different stages of 2D cortical differentiation showed that the targeted knock-in at the C-terminus of DCX in h9 hESCs is functionally correct. Thus, by differentiating the FnCas9-mediated endogenously tagged h9 hESCs towards 2D cortical neurons, we were able to visualize DCX-tdTomato expression and distinguish neuroblasts and early neurons from other cell types in the culture.

**Figure 2:**
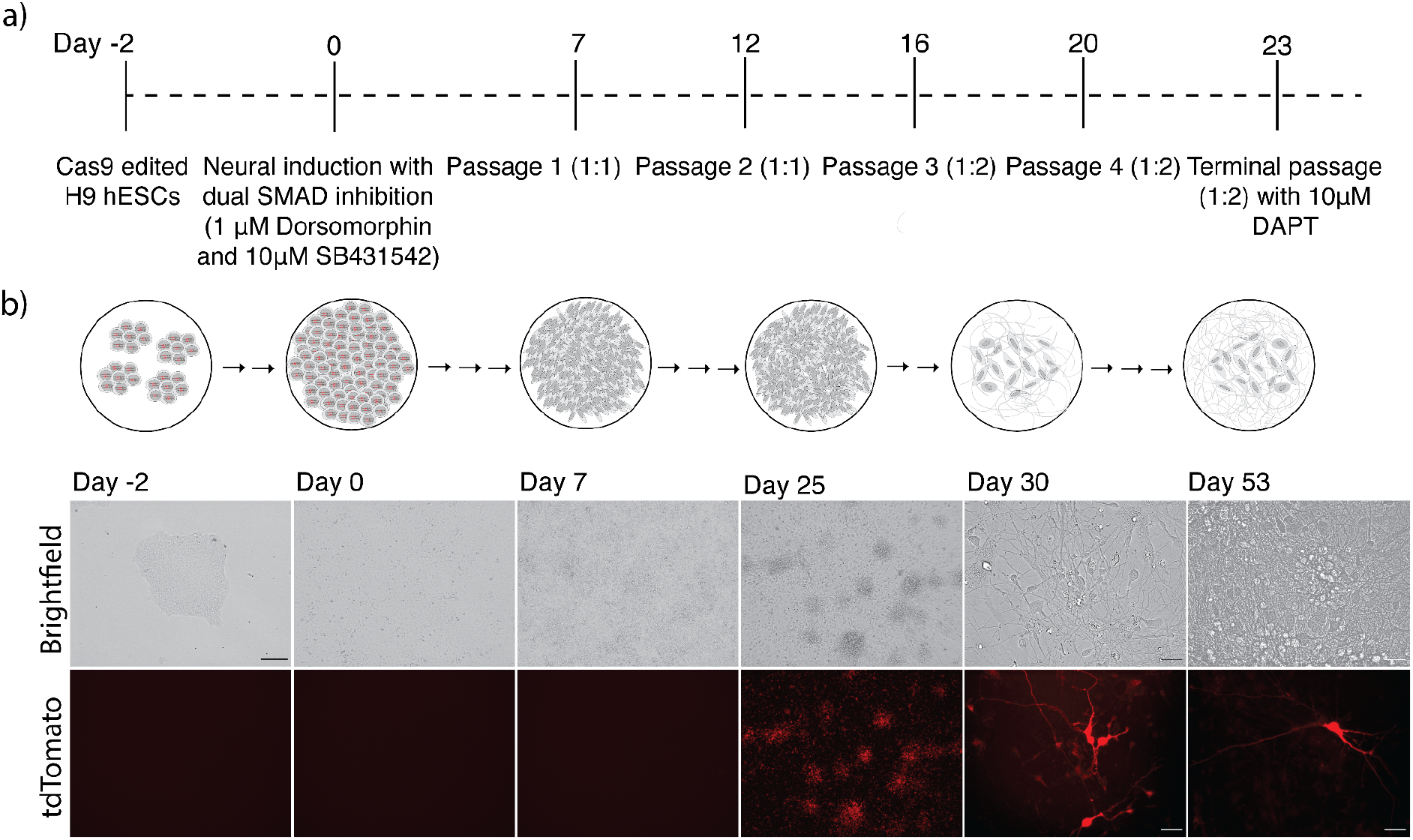
Stage-specific expression of DCX-tdTomato-marked neuroblasts in hESC-differentiated 2D cortical cultures; **a)** shows the optimized protocol for 2D cortical neural induction and expansion using a dual-SMADi approach that has been reported before (details are in the “Methods” section). **b)** Emergence of DCX-tdTomato as the cells start to express DCX, marking the presence of neuroblast subtypes between Days 25–30 and later immature neurons at Day 53 of the protocol. (scale: 50 µm for days 0 to 25; 10 µm for days 30 to 53)

Similar to DCX-tdTomato-tagged hESCs, TBR2-tdTomato-tagged hESCs also underwent differentiation, with expression evident as early as day 21 of the protocol (Supp. Figure 2). In summary, our results suggest that endogenous tagging of markers in iPSCs can allow the visualization of a subset of cells that express a desired gene. This is particularly important as the expression is fine-tuned to occur during a specific time point of 2D differentiation into the neural lineage.

### Stage-specific neural lineage tracing and purification in 3D cortical organoids

We next attempted to demonstrate the stage-specific expression of DCX-tdTomato in a more physiologically relevant 3D model, human cortical organoids (hCOs) (70, 71). Cortical spheroids and organoids represent a very useful model for studying various aspects of development and disease. Starting from human iPSCs, such cultures partially mimic certain aspects of the human brain that 2D cultures cannot efficiently model due to the lack of other cellular subtypes. Naturally, the possibility of dissecting specific neural lineages in a more complex 3D organ model is an attractive proposition for researchers investigating neurodevelopment with a high degree of granularity (71–75). Currently, single-cell RNA sequencing offers a snapshot of different cell types in organoids (71, 75), but such studies cannot be done without isolating cells by dissociating an organoid, a procedure that cannot be used for in-toto studies in growing organoid cultures.

We first tested h9 hESCs for cortical organoid (hCO) formation using a modification of a previously published protocol (Figure 3a, Methods) (76). Over the course of 75 days, these organoids were monitored for any observable or growth-related defects (Supp. Figures 3a and 3b). The neural rosette-like structures within these cortical organoids were visible as early as day 17 and became more distinct on days 24 and 33 (Supp. Figure 3a) (76,77).

**Figure 3:**
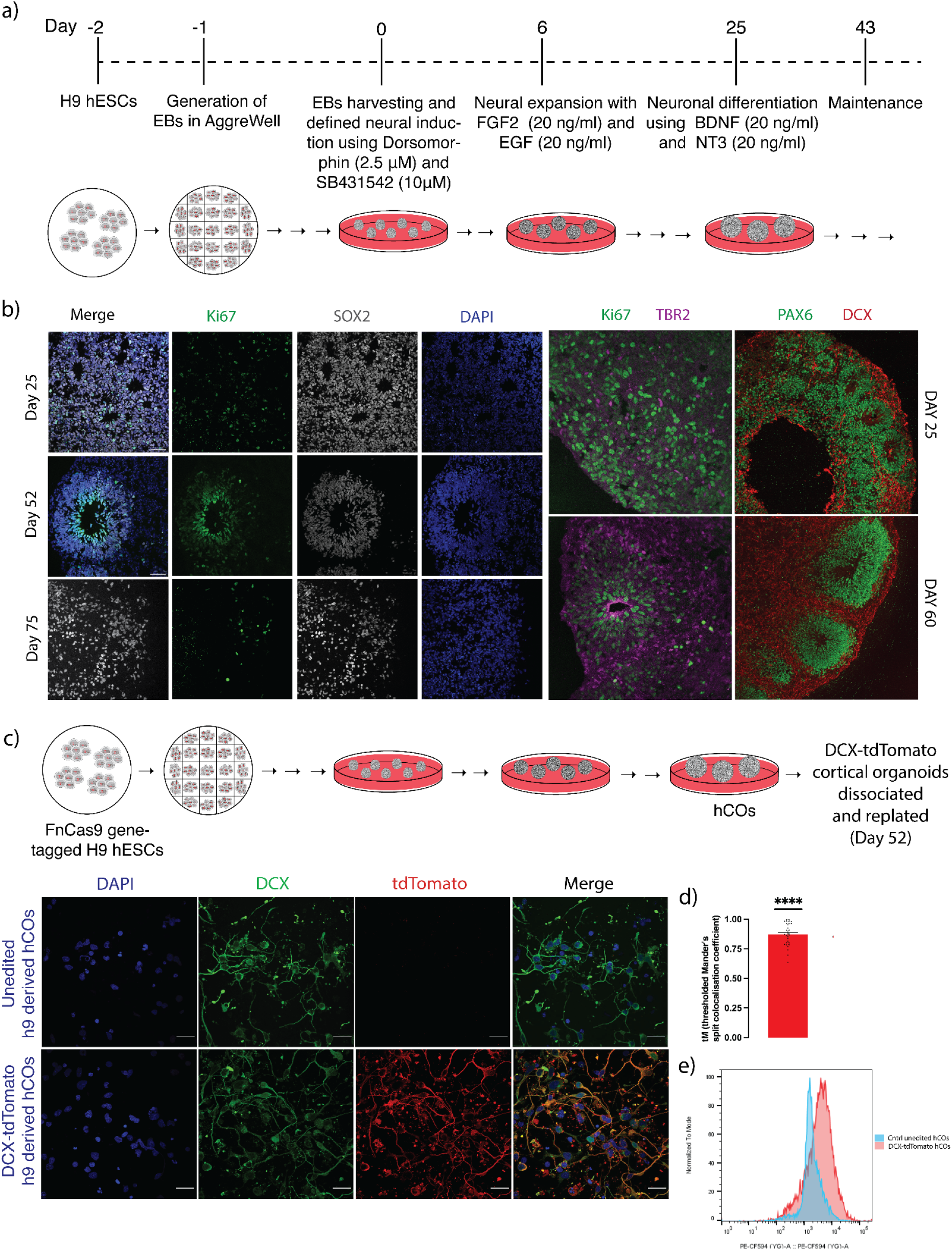
Generation of 3D cortical organoids (hCOs) from FnCas9-tagged hESCs and validation of the DCX-tdTomato expression. **a)** Schematic of the protocol optimized for the development of cortical organoids from the FnCas9-tagged hESCs (details in the “Methods” section). **b)** Left panel: expression of Ki67 and SOX2, highlighting the neural rosette cells and different developmental stages of cortical organoids (hCOs) at days 25, 52, and 75. Right panel: co-immunostaining of the organoid sections with neural marker proteins at days 25 and 60 showing their maturation (scale: 50 µm) **c)** Immunofluorescence-based co-localization of DCX and tdTomato expression within the cells dissociated from DCX-tdTomato-hCOs. (scale: 25 µm) **d)** The colocalization between the signals obtained from Red: Green channels are quantified as tM (thresholded Mander’s split colocalization coefficient), n = 30, error bars represent S.E.M., a single-mean two-tailed unpaired t-test is applied, and p values of **** ≤ 0.0001 (values from independent measurements are represented as dots). **e)** Real-time FACS-based cell sorting of live cells from DCX-tdTomato-tagged hCOs and unedited hCOs at day 52 (representative of three independent experiments).

Next, as shown in Supplemental Figure 3c, qPCR for early neurogenesis markers was done on RNA extracted from samples taken at different stages, such as h9 hESCs (H9), embryoid bodies (EB), days 25, 52, and 75, to look for stage-specific RNA expression during the different steps of the protocol. As can be seen, stemness markers such as NANOG and OCT-4 are only detectable at the h9, hESC, and EB stages, and these disappear with neural differentiation as expected (78–80). Also, SOX-2 and MYCN are highly expressed in h9 hESCs but are expressed even after neural differentiation by some neural subtypes such as neural stem cells, intermediate progenitor cells, etc., as seen on days 25, 52, and 75 (81–83). Expectedly, the expression of different neural lineage markers (PAX-6, ASCL1, TUJ1, TBR2, DCX, and TBR1) increased after neural induction (Supp. Figure 3c) (84-93). Overall, the stage-specific RNA expression profiling shows that the hESC-derived cortical organoids (hCOs) have different neural cell types that only show up at certain stages. Also, the expression of individual marker genes, such as PAX6, TBR2, DCX, and TBR1 could be mapped towards the maturation of these organoids (76, 77).

Interestingly, the expression of KI67, a proliferation marker, showed an increase on day 25 and a decrease on days 52 and 75, suggesting that the cells in hCOs can undergo neuronal mitotic and post-mitotic cell differentiation (Supp. Figure 3d) (89, 94). This also validated that the differentiation protocol was optimal since KI67 expression peaked at day 25 after FGF2 and EGF had caused neuronal expansion and went down as the hCOs matured, similar to human cortical development (76, 77). We confirmed this pattern of expression by co-labeling KI67 and SOX2 on cryosections of hCOs at days 25, 52, and 75. Also, through SOX2 and KI67 labeling, we observed neuroepithelial cells and neural stem cells at day 25, more dense neural rosettes at day 52, and a sharp decline in expression at day 75 (Figure 3b, left panel). Together, we could successfully monitor different stages in hCOs growth, such as neural induction (SOX2, PAX6), neural expansion (ASCL1, TBR2, TUJ1, DCX), and then neuronal differentiation (DCX, TBR1), in our 3D models (76, 77).

Next, we stained sections of these cortical organoids (hCOs) for Ki67, TBR2, PAX6, and DCX. At day 60, the rapidly dividing Ki67-positive cells were found to be lower than at day 25, while the differentiated TBR2-positive intermediate progenitor cells were higher. This showed the successful maturation of neurons in the cortical organoids (Figure 3b, right panel). A similar trend of continued maturation is seen at day 60 when DCX-positive neuroblasts or neuronal cells are more than PAX6-positive glia and NSCs (Figure 3b, right panel). Thus, stage-specific expression of markers across different stages of cortical organoids (hCOs) derived from unedited h9 hESCs demonstrates the efficacy of the protocol adopted for the growth of hCOs.

Once the growth conditions for cortical organoids derived from unedited h9 hESCs were well optimized and replicated, a similar protocol was used to create organoids from FnCas9-tagged DCX-tdTomato h9 hESCs. Cortical organoids derived from DCX-tdTomato h9 hESCs were once again monitored for growth and expansion. As shown in Supp. Figure 4a, the expression of different neural proteins like TBR2, TBR1, PAX6, and CTIP2 at day 52 could validate the neural identity of these cortical organoids. Also, the fact that these different neural proteins are expressed in hCOs made from DCX-tdTomato h9 hESCs shows that these cells don’t have any major disadvantages compared to h9 hESC-derived COs that haven’t been edited.

On day 52, after an optimal fluorescence signal, cortical organoids made from DCX-tdTomato h9 hESCs were dissociated and replated on laminin-coated dishes for immunofluorescence-based co-localization of the signal found by probing with anti-DCX (488 nm) and anti-tdTomato antibodies (594 nm) (Figure 3C). The cells from DCX-tdTomato hCOs show that tdTomato is being stably expressed, while the control cells from unedited hCOs show no endogenous expression of tdTomato (Figure 3C). Importantly, the endogenous tdTomato signal and antibody-labeled DCX showed very strong colocalization (Mander’s split colocalization coefficient of 0.8725; the details are in the methods section) (95, 96), which suggests that the endogenous marker is being expressed correctly in these 3D cultures. We used live DCX-tdTomato hCO cells for FACS to show that endogenously tagged DCX-tdTomato could be used to separate specific cell types. We could differentiate between cells from FnCas9-endogenously tagged DCX-tdTomato h9-derived hCOs and unedited hCOs (Figure 3e and Supp. Figure 4b) across their development. Finally, by immunostaining these sorted cells again with anti-DCX and anti-tdTomato, we were able to demonstrate that the targeted endogenous knock-in of tdTomato using FnCas9 at the DCX C-terminus could result in the efficient sorting of live DCX-positive cells from a heterogeneous population of cortical organoids (Supp. Figure 4c). Taken together, the use of FnCas9 to insert single-large DNA templates can be helpful for generating endogenously labeled cell types in both 2D and 3D models of neurodevelopment.

## Discussion

To study the different types of brain cells, it is important to be able to track the different neural lineages and do their molecular dissection. Even though there have been many other ways to do this in the past, they often can’t be used for efficient subtype tracing or purification in real time because of problems like non-random integration with viral-based methods or the need to destroy samples to get RNA or DNA for sequencing-based methods (97–99). An alternative method for tracking and isolating a neuronal subtype is through the endogenous expression of marker genes for marker-specific neural subtype tracing (100). The evolution of CRISPR/Cas9 systems for HDR-mediated endogenous gene knock-in has enabled us to do many of these things, such as *in vivo* gene tagging and real-time protein monitoring, etc. (46, 52–57). However, not all Cas9 systems effectively carry out such large HDR-mediated knock-ins. In a previous study, we found that Cas9 from the bacteria *Francisella novicida*, also called FnCas9, works better for HDR-mediated endogenous gene knock-in than canonical SpCas9 and its other high-fidelity variants (60). Using FnCas9’s ability to integrate single-large DNA templates efficiently, we were able to tag genes at their C-terminus in a single attempt endogenously. This was accomplished by using a DNA template as long as 4.1 kb that carried a fluorophore and an antibiotic resistance gene expressing through an independent promoter and floxed with loxP sites to be integrated at the C-termini of DCX and TBR2 within the genome of h9 hESCs, respectively. The successful integration at both DCX and TBR2 loci could then be confirmed by turning these endogenously marked h9 hESCs into cortical neurons and looking at the expression of tdTomato driven by endogenous DCX and TBR2 promoters. With the expression of tdTomato, the DCX-tdTomato h9 hESCs could also be visually distinguished from the other cells in the 2D cortical cultures. We then differentiated DCX-tdTomato h9 hESCs into 3D cortical organoids (hCOs) to show that tdTomato was only expressed in DCX-tdTomato h9-derived hCOs, which is a more physiologically relevant model. Furthermore, when tdTomato co-localization with DCX was evaluated in vivo, it was found that they coexist within neuroblasts, or early neuronal cells. Lastly, we were able to use DCX-tdTomato to use FACS to separate live DCX-positive cells from the different types of cells found in cortical organoids (hCOs).

In this article, we detail the development of donor constructs with distinct fluorescent tags and the use of FnCas9 to efficiently carry out HDR-mediated knock-in using a single, large DNA template. Two-dimensional (2D) cortical differentiation of hESCs that were genetically marked with an endogenous marker-specific fluorescent tag showed that an endogenous marker-specific fluorescent tag can be used to track IPCs (TBR2) and neuroblasts or immature neurons (DCX). By adding a fluorescent tag to the DCX locus, possible neuroblast cells in the hCOs could be traced. Also, using fluorescence-assisted cell sorting (FACS), it was possible to isolate these DCX-positive neurons. Although creating hESC models with multiple different colored fluorophores is laborious and time-consuming, it allows for the tracking of various cell lineages and the study of their interactions. On the other hand, the different colored fluorophores can be used to tag a single endogenous marker gene in distinct hESC populations, and they can be combined in varying proportions to create 2D or 3D cultures.

This kind of endogenous marker-specific gene tagging in hESC-derived cortical organoids can show the lineage transition more clearly than mouse analogues and can be used to study organ development step by step. Also, because these hCOs are grown in dishes, they can be used for live imaging by using marker-specific spatial labeling or FACS, to extract subtypes and study them in depth. In addition, the tools developed in this study can be applied to *in vivo* gene tagging using a single DNA template. But there are also some problems with using organoids for subtype tracing, such as their limited multiplexability, which requires maintenance and takes time, and the fact that not all cellular subtypes are present or come very late during maturation, such as microglia and astrocytes, and that their growth depends on the local environment and varies from batch to batch. Overall, this study shows that efficient FnCas9-mediated endogenous marker-specific fluorescent gene tagging can be a reliable way to track neural subtypes and pull them out of 2D or 3D models of neural cortical differentiation.

## Materials and Methods

### Plasmids and oligos used in this study

The plasmid with FnCas9 that was used in this was made and reported by our group (59). All the oligos used in this are listed in Supp. Table 1, along with a figure-wise mention of their use.

### Maintenance of h9 hESCs

The obtained h9 hESCs were maintained at 37°C and 5% CO_2_ in Essential 8 or E8 (Thermo Fisher Scientific) feeder-free maintenance medium with regular passages using Accutase (Thermo Fisher Scientific) and replated at optimal confluency on Geltrex-coated plates (Corning) with 10 µM Y-27632 ROCK inhibitor (STEMCELL Technologies) for the first 24 hours, followed by a media change to complete media with 100 U/ml penicillin/streptomycin (Thermo Fisher Scientific).

### FnCas9-based HDR-mediated endogenous gene-tagging in h9 hESCs

The plasmid containing FnCas9 Addgene 130969 (59) was incorporated with a crRNA sequence targeting the DCX C-terminus at the gRNA scaffold and was confirmed with PCR genotyping as well as Sanger sequencing. Following expansion, wild-type h9 hESC cells were supplemented with a 10 µM ROCK inhibitor (Y-27632, STEMCELL Technologies) 1 hour before nucleofection. The single cell suspension of 2×10^5^ h9 hESCs prepared through Accustase (Thermo Fisher Scientific) treatment was nucleofected using an Amaxa 4D nucleofector (CB-150 program) with a cocktail of 4 µg of a single plasmid containing FnCas9 + sgRNA and another 1 µg linearized donor DNA template in a P3 buffer (provided by the manufacturer). The nucleofected cells were immediately plated on a Geltrex-coated 6-well plate containing pre-warmed Essential 8 media (Thermo Fisher Scientific) containing ROCK inhibitor (Y-27632, 1:1000 to a final concentration of 10 µM) (STEMCELL Technologies) and Pen-Strep (1:100, Thermo Fisher Scientific) and maintained at 37°C and 5% CO_2_. The medium was changed to fresh Essential 8 for 24 hours after nucleofection. Further, after the next 24 hrs, the nucleofected h9 hESCs were selected with puromycin (0.4 µg/ml, selected by performing a minimum inhibitory concentration or MIC assay) for 48 hrs. Once the selected h9 hESCs grew back into colonies, their genomic DNA was taken out so that PCR genotyping could be done on the target locus.

### 2D cortical differentiation of genetically tagged h9 hESCs

FnCas9-tagged h9 hESCs were differentiated into cortical neurons in 2D cultures using a modified dual SMAD inhibition protocol from a previous report (69). First, optimally confluent hESCs were washed twice with Dulbecco’s phosphate-buffered saline (DPBS, Thermo Fisher Scientific), dissociated by ReLeSR (STEMCELL Technologies), and replated over the Geltrex-coated plates in Essential 8 medium (Thermo Fisher Scientific) and maintained at 37°C and 5% CO_2_, making sure to reach 100% confluency within 24 hours. After 24 hours on Day 0, a culture confluency of 80–100% is ideal for neural induction. (Wait another day before introducing the neural induction if the confluency is not 80–100%.) For the neural induction, the medium was then replaced with N2/B27 medium (1x N2 supplement, 1x B27 minus Vit. A supplement, and 1x Glutamax in Neurobasal-A media) containing the SMAD inhibitors, 1 µM dorsomorphin (STEMCELL Technologies), and 10 µM SB-431542 (STEMCELL Technologies), and maintained at 37°C and 5% CO_2_. The neural induction was performed by replacing N2/B27 (+SMADi) daily for 7 days. On the 7^th^ day, to perform passage 1, the multi-layered neuroepithelial cells were washed with Hanks’ Balanced Salt Solution (HBSS, Thermo Fisher Scientific), mildly dislodged as clumps using Accutase (Thermo Fisher Scientific), and replated over freshly coated Geltrex plates using N2/B27 medium (SMADi) containing 10 µM Y-27632 ROCK inhibitor (STEMCELL Technologies). With the ROCK inhibitor removed after 24 h (day 8), just the N2/B27 medium without SMADi was replaced daily until day 12. On day 12, adhered neural progenitors were again dislodged using Accutase (Thermo Fisher Scientific) and replated over Geltrex-coated plates within N2/B27 medium containing 10 µM Y-27632 ROCK inhibitor (STEMCELL Technologies). After the removal of 10 uM Y-27632 ROCK inhibitor on day 13, cells were maintained at 37°C and 5% CO_2_ until day 23, with daily media changes using just the N2/B27 media and in-between passages (1:2) on days 15–16 and 19–20 as per previous. On Day 23, to direct the terminal neuronal differentiation, the cells were passaged (1:2) in a B27-only medium (1x B27 minus Vit. A supplement and 1x Glutamax in Neurobasal-A media) supplemented with 10 μM DAPT (Sigma-Aldrich) and 10 µM Y-27632 ROCK inhibitor over poly-l-ornithine + laminin-coated plates and left till days 25–26 with no media changes at 37°C and 5% CO_2_. On the 26^th^ day, half of the media was replaced with half the initial volume of fresh B27+DAPT. From days 29–30 on, differentiated neurons were maintained in the B27-only medium with half-volume media changes once a week or when necessary.

### Generation of h9 hESC-derived cortical organoids (hCOs)

A published protocol from Paşca, Anca M., et al. (2015) (76) was a bit modified to turn unedited or FnCas9-tagged hESCs into cortical organoids (hCOs). The h9 hESCs were grown on a Geltrex-coated plate in Essential 8 medium (Thermo Fisher Scientific) at 37°C and 5% CO_2_. When the cultures reached 70–80% confluency (every N5 day), they were replated using ReLeSR (STEMCELL Technologies) into E8 medium with ROCK inhibitor (Y-27632, 1:1000 to a final concentration of 10 μM) (STEMCELL Technologies). On day −2, the ROCK inhibitor is removed and the medium is replaced to complete E8. On day −1, a single cell suspension of the cultured hESCs using Accutase (Thermo Fisher Scientific) was prepared, strained through a 40 µm filter, and transferred to a low attachment Aggrewell plate (STEMCELL Technologies), achieving a final concentration of 3×10^6^–4×10^6^ cells per well, or 10,000–12,000 cells/embryoid body (EB) in E8 medium with ROCK inhibitor. Once evenly distributed, the cells were kept undisturbed at 37°C and 5% CO_2_ for the next 24 hours to form embryoid bodies. The neuralization began on day 0 by transferring the EBs to ultra-low attachment 6 cm dishes (Corning) and adding fresh Essential 6 or E6 medium (Thermo Fisher Scientific) supplemented with dorsomorphin (DM; 2.5 µM final dissolved in DMSO) (STEMCELL Technologies) and SB431542 (LDN; 10 µM final dissolved in ethanol) (STEMCELL Technologies). Media changes (skipping day 1, 24 h) were performed daily for the first five days with fresh E6 medium containing DM and LDN. On the sixth day in suspension, the floating organoids in the same ultra-low attachment plates were replaced with neural medium (NM) that had Neurobasal-A (Thermo Fisher Scientific), 1x B-27 (Thermo Fisher Scientific) minus vitamin A, 1x GlutaMax (1:100), and 1x Pen-Strep (1:100), and were maintained at 37°C and 5% CO_2_. The NM was supplemented with 20 ng/ml FGF2 (Gibco) and 20 ng/ml EGF (Gibco) for the next 19 days, with daily media changes for the first 10 days and every alternate day for the subsequent 9 days. Once EGF and FGF2 are added to the medium, the organoids will grow quickly. When needed, they should be moved to 10-cm ultra-low attachment dishes (Corning), making sure that after each change of media, the cortical organoids are spread out on the plate and not all in the middle. They should then be put back in the incubator, maintaining 37°C and 5% CO_2_. Last, FGF2 and EGF were replaced with 20 ng/ml BDNF (Gibco) and 20 ng/ml NT3 (Gibco) on day 25 to help the neural progenitors turn into neurons. From day 43 on, only NM without growth factors was used for every four-day media change. Organoids were monitored for their growth and accordingly distributed into other 10-cm dishes to ensure that the medium pH remained stable.

### qPCR-based neural marker expression validation of the hCOs

The RLT lysis buffer (Qiagen) was used to break up cortical organoids that were taken at different times. Following the RNeasy Micro Kit (Qiagen) instructions allowed for the extraction of RNA. To get rid of genomic DNA, a 15-minute on-column digestion with RNase-free DNase I (Qiagen) was added. Then, using a High-Capacity cDNA Reverse Transcription Kit (Thermo Fisher Scientific) and the manufacturer’s instructions, the purified RNA was turned into cDNA. To do qRT-PCR, reactions were made with TaqMan Universal PCR Master Mix (Thermo Fisher Scientific) and different commercially available TaqMan probes (Supp. Table 1) according to the manufacturer’s instructions. The reactions were then run in triplicate on the Lightcycler 480 II thermal cycler (Roche, Switzerland). The −2-DDCt or Livak method (101) was used to analyze the data, and relative expressions normalized against HPRT1 were plotted using GraphPad Prism.

### Cryo-sectioning and immunofluorescence of the hCOs

hESC-derived cortical organoids at different developmental stages were taken out and washed three times with Dulbecco’s phosphate-buffered saline (DPBS, Thermo Fisher Scientific). Once thoroughly washed, they were fixed for 20 to 30 minutes with 4% paraformaldehyde (PFA) and washed three times again with DPBS before being kept overnight in 30% sucrose. After their complete immersion in sucrose, the organoids were embedded in a 1:1 OCT-tissue embedding medium (Leica):30% sucrose solution and stored frozen at −80°C. The cryo-blocks containing cortical organoids were cryo-sectioned at 10 µm thickness, and the sections were collected on SuperFrost slides (Thermo Fisher Scientific) to preserve at minus 20°C until performing immunohistochemistry. The sections were allowed to dry for at least 10–20 minutes at RT, and the sectioned tissue area was outlined using a hydrophobic PAP pen (Thermo Fisher Scientific). The cryosections were then rehydrated to remove excess OCT. Following three washes in PBS for 5 minutes, the sections were once again post-fixed directly on the slides using 4% PFA for 10 minutes at room temperature (RT). They were then blocked for 1 hour at RT using a blocking solution (0.3% Triton X-100 in PBS plus 10% normal donkey serum). Overnight incubation at 4°C with the primary antibody (Supp. Table 2) diluted in blocking solution was performed. After being washed three times with PBS, sections were put in a blocking solution with Cy3, Alexa Fluor 488, and 647 labeled secondary antibodies (1:500) from Jackson’s laboratory for an hour at room temperature. Finally, the sections were rewashed three times and counter-stained with DAPI (1:10,000; Sigma-Aldrich) to be mounted using VECTASHIELD Antifade Mounting Medium (Vector Laboratories) and stored at 4°C until imaging.

### Dissociation of hCOs followed by immunofluorescence

Cortical organoids were washed three times in DPBS and put in Accutase (Thermo Fisher Scientific) for 20 minutes at 37°C to make 2D neuronal cultures that could be used for immunofluorescence-based colocalization. hCOs were then triturated into a single-cell suspension by gently pipetting up and down using a p200 tip. The resulting suspension was filtered through a 40 µm filter, centrifuged at 300 g for 5 minutes, and plated onto Geltrex-coated coverslips in a 24-well plate with neural medium (NM) containing Neurobasal-A (Thermo Fisher Scientific) supplemented with B-27 supplement without vitamin A (Thermo Fisher Scientific), GlutaMax (1:100, Thermo Fisher Scientific), and Pen-Strep (1x, Thermo Fisher Scientific). After 24 hours, the adhered cells were washed three times with DPBS and fixed using 4% paraformaldehyde (PFA). Following fixation, the cells were washed three times with DPBS and blocked for an hour at room temperature with a blocking solution (0.2% Triton X-100 in PBS plus 10% normal donkey serum). The cells were incubated overnight at 4 °C with diluted primary antibodies (Supp. Table 2). After three washes with PBS, the cells were left at room temperature for an hour with a secondary antibody diluted in blocking solution (1:500) with Alexa Fluor 488 or 568 labeled secondary antibodies from Invitrogen-Molecular Probes. Finally, after the 3x PBS washes, the cells were counterstained with DAPI (1:10,000; Sigma-Aldrich), and the slide was mounted using VECTASHIELD Antifade Mounting Medium (Vector Laboratories) to be stored at 4°C until imaging.

### Image acquisition and co-localization

The images were acquired using a laser-scanning TCS SP5 II confocal microscope (Leica Microsystems) at a z-section thickness of 1 µm. For image analysis and visualization, the software FIJI ImageJ was used (102). For the co-localization signal quantification, the maximum-intensity Z-projected images were run through an ImageJ plugin called Colocalization Threshold. This gave the thresholded Mander’s split co-localization coefficient, and the values were plotted using GraphPad Prism software.

### Dissociation of hCOs followed by fluorescence-activated cell sorting (FACS)

Cortical organoids were washed three times with DPBS and treated with Accutase (Thermo Fisher Scientific) for 20 minutes at 37°C to get 2D neuronal cells for the FACS. The hCOs were then gently pipetted up and down using a p200 tip. The resulting cell suspension was filtered through a 40 µm filter and centrifuged at 300 g for 5 minutes. The cell pellet was resuspended in DPBS and transferred into the FACS tube to perform FACS using a cell sorter (BD FACSMelody, BD Biosciences). Live events during FACS were accordingly recorded and plotted using the software FlowJo.

### Statistics

The data are presented as the mean ± standard error of the mean (SEM), and a single mean unpaired two-tailed t-test was used to determine statistical significance (GraphPad Prism). Details of the statistical analyses are found in the figure legends.

## Acknowledgement

The authors would like to acknowledge all members of Dr. Debojyoti Chakraborty and Dr. Souvik Maiti’s lab for their insightful discussions and contributions to this study. We’d also like to thank the members of the Neural Stem Cell Biology Lab at the Francis Crick Institute in the UK for their helpful ideas and for hosting the cortical organoids-related experiments. We would like to thank the Human Embryo Stem Cell Unit (HESCU) at the Francis Crick Institute, as well as the confocal imaging facilities at the Francis Crick Institute and CSIR-IGIB, especially Dr. Himanshi Kapoor. The authors would also like to thank Ms. Antara Sengupta and Ms. Riya Rauthan for providing helpful input during the execution of experiments for this study. Through the Newton Bhabha Fund PhD Placements Programme 2018–2019, which M.K. received, and the Department of Biotechnology (DBT) grant GAP188 to D.C., the British Council in India and the Department of Biotechnology (DBT) partially funded this study.

## Author’s contribution

M.K. conceived the project and designed the experimental pipeline with inputs from D.C. and F.G. M.K. and B.T.T. performed hCOs validation experiments. M.K. drafted the manuscript with input from D.C. and others.

## Ethics declarations

The authors declare no competing interests pertaining to this work.

## Notes

### Competing Interest Statement

The authors have declared no competing interest.

