## Supplementary Figures for "Endogenous gene tagging with FnCas9 to track and sort neural lineages from 3D cortical organoids"

a) FnCas9-mediated HDR-based locus-specific knock-in for endogenous DCX-tdTomato validated by Sanger sequencing.

DCX C-teminal knock-in forward direction

DCX C-teminal knock-in reverse direction

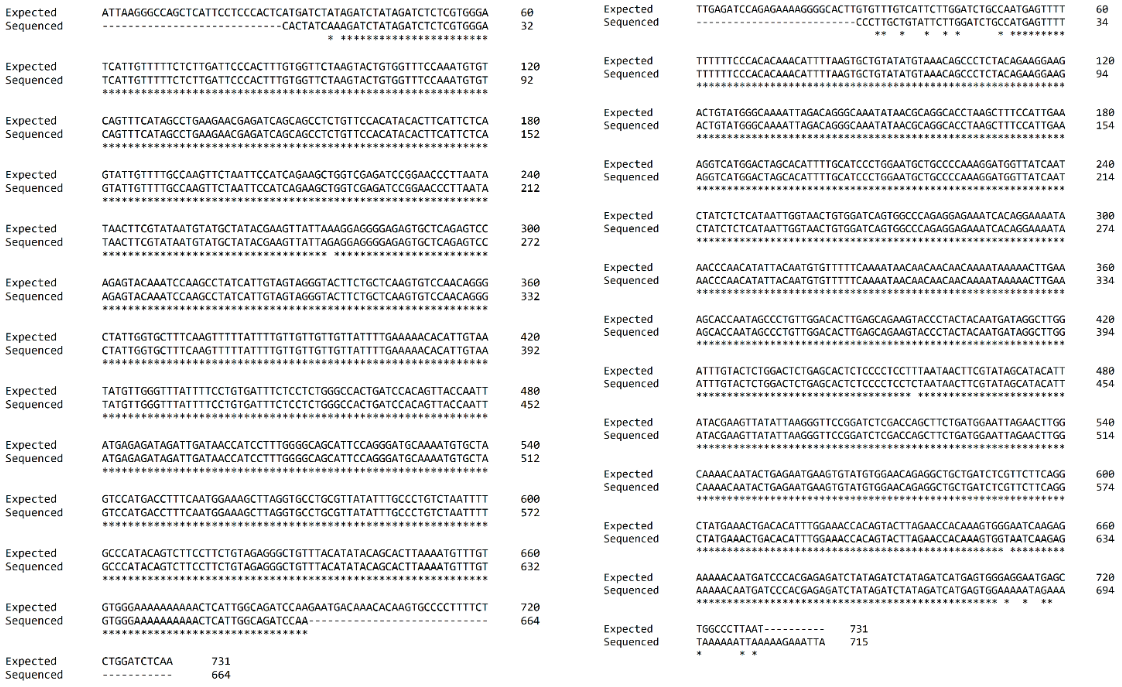

b) HDR-based locus-specific knock-in on the endogenous Tbr2 C-terminus using FnCas9.

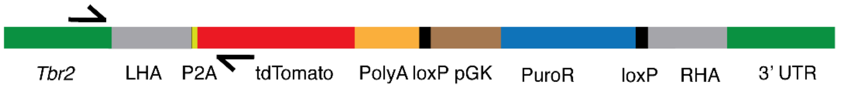


Tbr2 C-terminal locus after sucessful integration of the dsDNA template

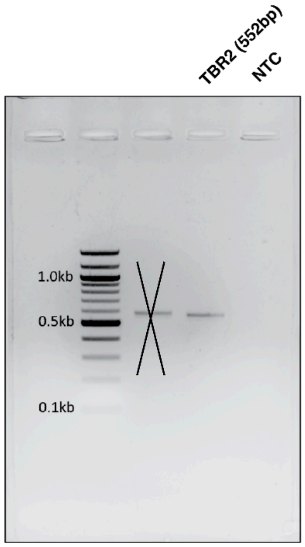

**Supp. Figure 1, a)** Results of the Sanger sequencing from both forward and reverse directions at the C-terminus end of the *DCX-tdTomato* integration site **b)** The linearized donor DNA template used for knock-in with its various components, along with left and right homology arms complementary to the C-terminus end of the endogenous *TBR2.* A 552-bp DNA band on the agarose gel confirms the PCR genotyping of the antibiotic-selected cells with the successful integration of the DNA template used. (NTC, no DNA template control)

2D - neural differentiation to visualise the lineage-specific expression of fluorescently tagged endogenous marker gene Tbr2-tdTomato.

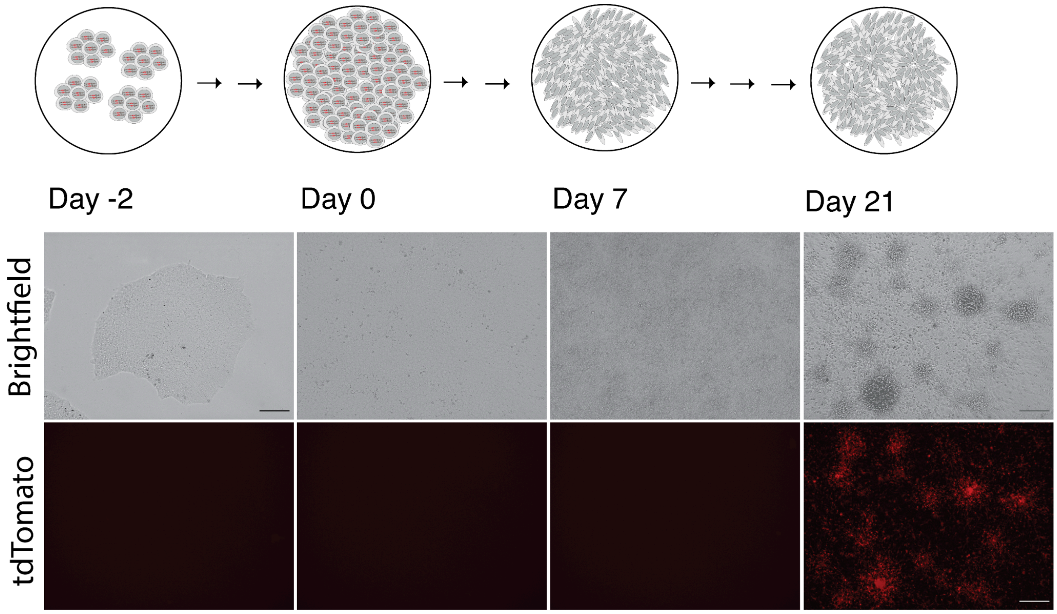


**Supp. Figure 2** shows the expression of TBR2-tdTomato and the presence of intermediate progenitor cells (IPCs) on Day 21 of the 2D neural differentiation protocol (scale: 50 μm).

Visual growth monitoring and expression validation of various neural genes within hCOs derived from H9 hESCs

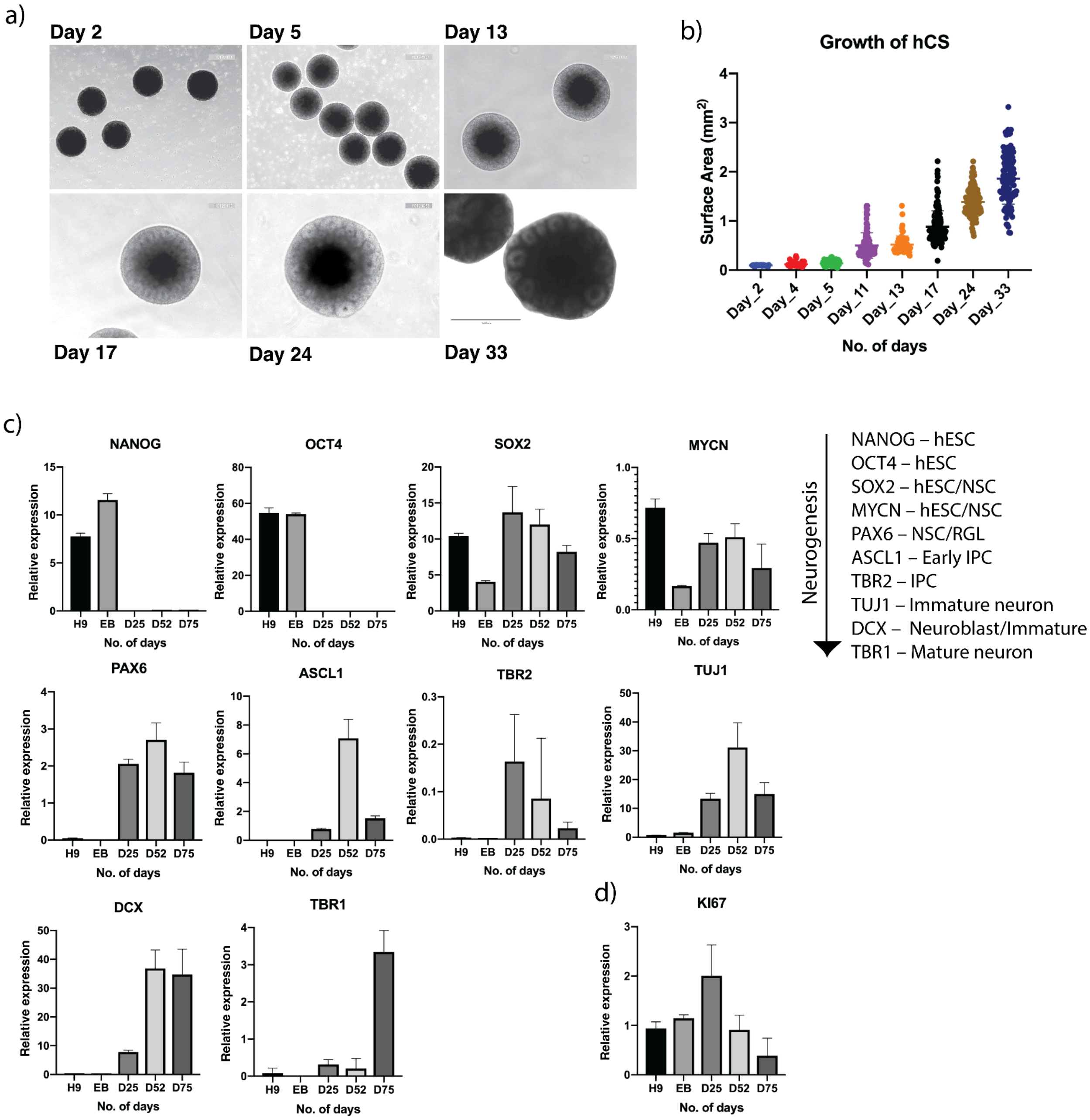


**Supp. Figure 3, a)** Day-wise monitoring of h9 hESC-derived cortical organoids (hCOs); visible neural rosettes could be seen starting from day 17 of the used protocol. (scale: 1OOO µm) **b)** A total surface area-based quantification shows steady growth of the cortical organoids with comparatively less heterogeneity in their size. **c)** The expression of various neurogenesis genes could confirm the progress of neural differentiation and maturation across the cortical organoids**.** (qRT-PCR analysis was performed on samples obtained from a minimum of three independent batches of COs.) **d)** The hCOs could also imitate neuronal mitotic and post-mitotic cell differentiation, with Ki67 expression peaking at day 25 after FGF2 and EGF-caused neuronal expansion.

The visual expression of various neural proteins on day 52 of the protocol in cells replated from dissociated cortical organoids grown from DCX-tdTomato tagged h9 hESCs.

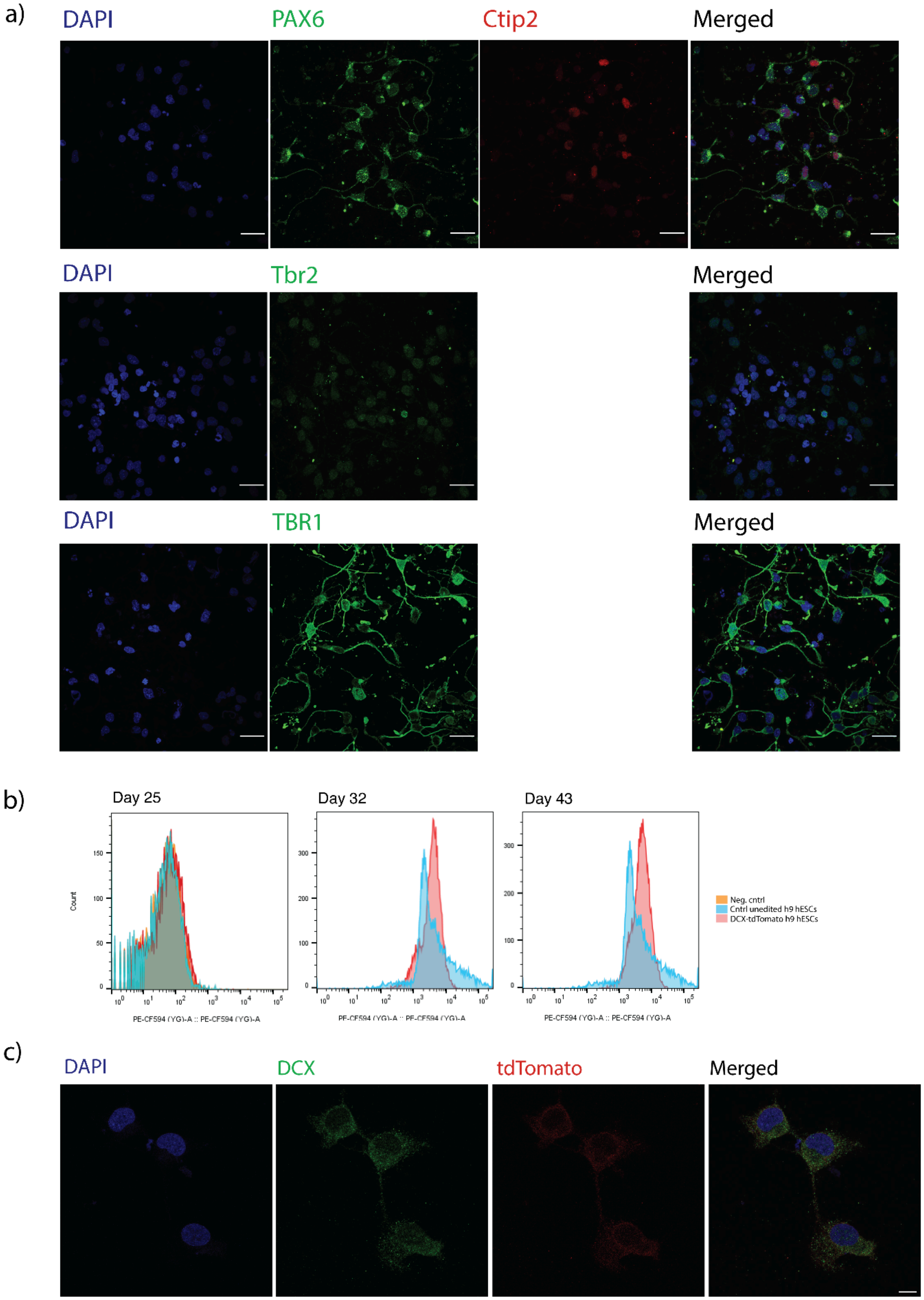

**Supp. Figure 4, a)** Expression of different neural proteins at day 52 of the protocol in the cells replated from dissociated cortical organoids derived from DCX-tdTomato-tagged h9 hESCs (scale: 25 μm) **b)** FACS-based detection of DCX-positive cells in real-time from the live hCOs at days 25, 32, and 43 (representative of three independent experiments). **c)** Immunofluorescence of the DCX-tdTomato-positive cells after FACS over the cortical organoids (scale: 10 μm).

### Supplementary tables:

- **Supp. Table 1,** List of oligos and probes used for this study
- **Supp. Table 2,** List of antibodies used for this study
